# Temporal span of biodiversity monitoring mediates the effects of area and environment

**DOI:** 10.64898/2026.02.09.704769

**Authors:** Daniela Mellado-Mansilla, Gabriele Midolo, Gabriel Ortega-Solís, Jiří Reif, Florencia Grattarola, Dylan Craven, François Leroy, Michela Perrone, Karel Šťastný, Vladimír Bejček, Petr Keil

## Abstract

The scale at which diversity is observed shapes the patterns we find. While spatial scale is known to influence biodiversity patterns, the effects of temporal scale, namely the average duration of sampling (known as temporal span), have been mostly overlooked. Here, we investigate how temporal span affects species richness patterns, their environmental drivers, and species richness hotspots. We used species richness data from several large bird datasets from Czechia, with over 7000 observations, a spatial grain ranging from 0.03 to 100 km^2^, and a temporal span ranging from 1 to 36 years (1985-2017). Using Random Forests, we modelled species richness as a response to temporal span, while also including area, geographic location, time, and environmental and land-cover predictors. We found that the temporal span is consistently among the most important predictors of bird species richness. Moreover, temporal span interacts with key environmental conditions, particularly precipitation and water bodies, modulating their effects on species richness and revealing processes that differ from those traditionally attributed solely to spatial grain. We also found that using different time spans can shift the predicted locations of biodiversity hotspots. Our results provide empirical evidence that temporal span should be included in studies about biodiversity and conservation planning, given the urgent challenges arising from ongoing biodiversity change and the complexity of its drivers.

## Introduction

Ecologists are interested in where biodiversity is, how it changes over time, and what drives it. For the simplest and most widely-used metric of biodiversity – species richness – well-established patterns include its latitudinal and altitudinal gradients (Ricklefs 2004, Lomolino et al. 2017). In parallel, climate, productivity, and habitat type have been identified as key environmental predictors of species richness (Field et al. 2009). Yet, biodiversity patterns and the effect of the environment on these patterns are strongly dependent on the *spatial grain* (i.e., the average area where species are recorded) (Rahbek 2005, Field et al. 2009, Belmaker and Jetz 2011). The most fundamental form of this grain dependence is the *species-area relationship* (SAR), i.e., the increase in the number of species with increasing area (Storch 2016). SAR emerges through sampling effects (Gooriah and Chase 2020), extinction and colonization (dispersal) dynamics (MacArthur and Wilson 1967), and environmental heterogeneity (Williams 1943). These mechanisms also explain the grain-dependence of spatial diversity patterns, specifically through geographic variation in species turnover, or beta diversity (Keil and Chase 2019). Accordingly, scale dependence has become a central topic of biodiversity research (Storch et al. 2007), and many recent analyses explicitly account for spatial grain to better capture the complexity of diversity patterns (Field et al. 2009, Belmaker and Jetz 2011, Keil and Chase 2019, Craven et al. 2020).

The *species-time relationship* (STR), which describes how the species observed at a given site accumulate over time (Preston 1960, Adler and Lauenroth 2003). When time is the measure of sampling effort in STR, it is often defined as the discovery curve or collector’s curve (Colwell and Coddington 1994). This is a special case of the general concept of species accumulation curves (Gotelli and Colwell 2001), which describe how species richness increases as a function of the sampling effort. Furthermore, the effects of time and space on diversity can be integrated into the *species-time-area relationship* (STAR; Adler & Laurenoth 2003, Adler et al. 2005), which captures how the parameters of a SAR can shift over time (Adler & Laurenoth 2003, Adler et al. 2005). As a result, the spatial scale-dependency of biodiversity should be affected strongly by the temporal span of observations, and thus accounting for the temporal span may also help explain why some studies sometimes reach conflicting and often counterintuitive conclusions about biodiversity patterns. Many results described as “context dependent” (Catford et al. 2022, Spake et al. 2023) in ecology could partly reflect differences in temporal span. For instance, studies on the relationship between biodiversity and ecosystem functioning often report high variability across sites (Ricklefs 2004), which may partially arise from variation in this relationship over time (Guerrero-Ramírez et al. 2017). By explicitly accounting for the temporal span when modeling biodiversity, we anticipate uncovering novel drivers of species diversity patterns.

Despite its potential importance, the effects of temporal span on diversity patterns and their environmental drivers have received little attention (but see White 2004, White et al. 2006, 2018, Erős and Schmera 2010, Swenson et al. 2013). Here, we provide a comprehensive empirical assessment of the effect of temporal span on patterns and drivers of species richness, using a dataset of long-term bird observations from Czechia of over 30 years at spatial scales ranging from 0.03 to 100 km^2^. Specifically, we (1) assess the relative importance of time span versus area and environmental drivers across different spatial scales and data types; (2) quantify the empirical STAR curves resulting from our models; (3) evaluate how time span modulates species-environment relationships; and (4) investigate how variation of time span affects the detection of hotspots of species richness predicted from the species-environment relationships.

## Methods

We quantified the number of bird species (species richness) across Czechia using three complementary data sources that span multiple spatial grains and time spans. We modeled species richness as a function of area, time span, and a suite of environmental variables, including climate, land cover, vegetation productivity, and protected area surface, using the Random Forest algorithm. We then examined the relative importance of temporal span and environmental predictors and assessed how time span modulates the effects of other predictors. We then predicted species richness using three different time spans.

### Bird data

We used three sources of data on Czech bird species richness (Figure 1a):

**Figure 1.**
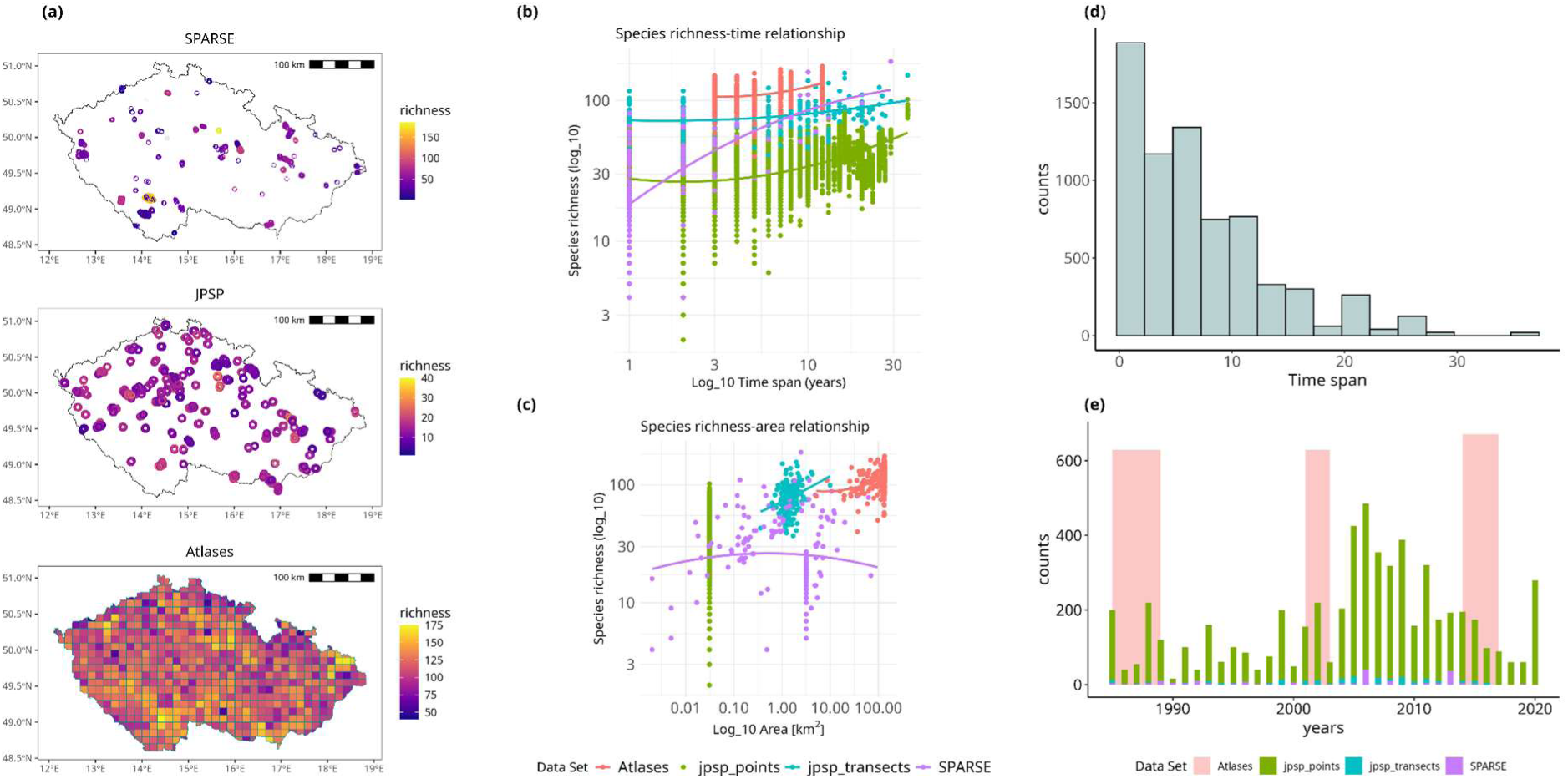
(a) Three data sets of bird occurrences across Czechia and their observed time-species richness relationships, time spans (b) and area-species (c) richness relationships (fitted with quadratic regression); (d) histogram indicating the number of sampling events per time span, and (e) the sampling events per year (as the start year). Maps show species richness for each data set at different periods: SPARSE, from 1980 to 2018; JPSP, from 2000 to 2010; and for Atlases, the species richness of each cell was calculated by randomizing the three different versions and their combinations (see Methods). Individual plots of the species-area and species-time relationships can be found in Supplementary Fig. 1 and Supplementary Fig. 2, respectively.

#### Standardized coarse grid atlases (Atlases)

First, we used three gridded breeding bird atlases (Šťastný et al. 1997, 2006, 2021) that include surveys conducted over three time periods: 1985-1989, 2001-2003, and 2014-2017. These atlases cover the entirety of Czechia with a spatial resolution of approximately 10 x 10 km² (Fig. 1), with around 630 cells surveyed repeatedly in each period.

#### Standardized local surveys (JPSP)

The second dataset that we used is the Common Bird Monitoring Program (Jednotný program sčítání ptáků, hereafter JPSP) (Reif et al. 2013), which spans from 1982 to 2020. This dataset consisted of 300 sampling transects (*JPSP Trs*), each containing 20 points (*JPSP Pts*) with a diameter of approximately 0.03 km^2^ and separated by 300-500 m. The transects are randomly distributed across Czechia and were consistently sampled each year using a fixed methodology (see Reif et al. 2013 for more details). For the rest of our analyses, JPSP points and transects were analyzed separately.

#### Heterogeneous inventories (SPARSE)

The third dataset was the SPARSE database (Tschernosterová et al. 2023), an open database that compiles information on bird species composition and richness from 1890 to 2018 with areas ranging from 0.002 to 95 km^2^, for 348 heterogeneous geographic units (i.e., polygons, transects, and points) in Czechia that uses different sampling methods. SPARSE collects data from published local inventories, reserve surveys, and fauna lists. SPARSE geographic sampling units are distributed across Czechia, and cover different types of habitats (Fig.1).

We used the area of each geographic unit (i.e., point, transect, polygon, or grid cell) in km^2^ as a predictor in all models.

### Survey and time span definitions

Each geographic unit in the three datasets was surveyed during one or more years. We define a *survey* as an uninterrupted period of consecutive years where we have data on species richness for each year. When there were multiple sampling events within one year (e.g., in May and then in July), we pooled them so that they represent one year. Years with no data on species richness break the continuity of a survey, separating it into two or more distinct surveys. For example, at a given geographic unit, there can be a 3-year survey conducted in 1987-1989 and a 5-year survey in 1991-1994. Each survey has its own species richness, and these surveys are then the fundamental observations in all our analyses. We define the *time span* of a survey as the number of consecutive years in the survey, i.e., its duration in years.

### Species richness of surveys

For the *SPARSE and JPSP* datasets, we selected geographic units with data from 1985 onwards (Fig. 1d & e) to ensure temporal overlap across all datasets. For each survey in each geographic unit, species richness was estimated as the number of unique species recorded. This gave us a total of 6445 surveys with a median time span of 6 years per survey (mean± SD = 7.2 ± 6.3 years, range = 1–36 years).

The *Atlases* dataset has three editions covering 5, 3, and 4 years, respectively; i.e., each grid cell has three surveys. This would give us a maximum time span of 5 years, which is much less than in the JPSP dataset. To generate time spans comparable to those in the SPARSE and JPSP datasets, we merged some of the atlas surveys even though they include time gaps while avoiding spatial pseudoreplication (i.e., the same cell appearing in multiple time spans). We addressed both problems by randomly assigning the 628 grid cells of each atlas into six non-overlapping groups (∼100 cells each; no cell shared across groups), each corresponding to a unique combination of atlases and sampling periods: (1) atlas 1 only (1985-1989, 5-year span), (2) atlas 2 only (2001-2003, 3-year span), (3) atlas 3 only (2014-2017, 4-year span), (4) atlas 1 and 2 combined (8-year span), (5) atlas 2 and 3 combined (7-year span), and (6) the three atlases combined (12-year span) (See Supplementary Fig. 3). The species richness for each cell was calculated by aggregating all unique species recorded during the time span covered by the atlases or the combination of atlases assigned to a given cell.

### Environmental predictor variables

The environmental predictors used in our models were selected based on the results of previous biogeographical studies linking the diversity of Czech birds with the environment (Storch et al. 2003, 2023, Moudrý and Šímová 2013, Prajzlerová et al. 2024).

#### Climatic predictors

We extracted data on mean annual temperature, minimum temperature, maximum temperature (°C), and precipitation (mm) at 1 km^2^ resolution from CHELSA (Karger et al. 2021) for each sampling unit. For each year a unit was surveyed, we calculated annual averages of the corresponding variables and then computed overall means across the total period sampled for each unit.

#### Land cover

We extracted land cover data from CORINE at a 100 x 100 m resolution for the years 1990, 2000, 2006, 2012, and 2018 (European Union’s Copernicus Land Monitoring Service information 1990, 2000, 2006, 2012, 2018). We used the level 1 of landcover classes which includes: 1) artificial surfaces, 2) agricultural areas, 3) forests and seminatural areas, 4) wetlands, and 5) water bodies. For each geographic unit, we overlaid the CORINE rasters and calculated the total area (in km²) occupied by each land cover class within its boundaries. Since CORINE 1990 recorded information from around 1985 onward, we used this version to represent land cover between 1985 and 1990.

#### NDVI

We used Google EarthEngine (Gorelick et al. 2017) to extract the Normalized Difference Vegetation Index (NDVI) at a 250 x 250 m resolution from MODIS (Didan et al. 2015), for June and July, which corresponds to the peak of the growing season. NDVI data are available from the year 2000 onward. Pixels were first masked by the quality layer of the MODIS dataset to minimize possible effects of cloud ice or values flagged as unreliable. Overlapping MODIS images were reduced to a single layer by creating a median-pixel composite. Then, we calculated the median NDVI during the time period in each geographic unit.

#### Protected areas

We also obtained data on the total surface area of protected areas (in km²) within each geographic unit. Specifically, we extracted the area of small-scale protected sites and large-scale landscape protected areas, as provided by the Nature Conservation Agency of the Czech Republic (Nature Conservation Agency of the Czech Republic 2025). For each sampling unit, we summed the areas of these two types of protected sites to calculate the total protected area.

### Random forest models

We fitted a Random Forest model (Breiman 2001) for each of the four data sets (Atlases, SPARSE, and JPSP points, and JPSP transects), with a total of 628, 172, 5,974, and 299 observations, respectively. Additionally, we fitted a model for all the data sets together. The response variable was species richness per geographic unit and sampling period. The predictor variables were the time span of the survey per sampling unit, the first year of the sampling period (i.e., start year), the area of the sampling units (in km^2^), the centroid latitude and longitude, and all the environmental variables described above. We included the latitude and longitude as predictors to account for spatial autocorrelation (following Viana et al., 2022), and the start year in a similar way to account for temporal autocorrelation. By doing so, this modeling framework accounts for spatio-temporal heterogeneity and also facilitates the integration of multiple data sources (Keil and Chase 2022).

To reduce multicollinearity in our models, we first estimated the correlation between all pairs of predictors. Then, we ran a Random Forest including all predictor variables for each dataset and extracted the variable importance scores. We examined all pairs of predictors with a Pearson coefficient higher than 0.6 and retained the variable with the highest importance scores. We then re-fitted our models without the excluded variables (see Supplementary Table 1 for details on each model’s final predictors). Each model was tuned by adjusting the number of trees from 500 to 3000, the minimum number of samples required per split from 2 to 5, and from 5 to 26 candidate predictors per split using 72 possible combinations in the hyperparameter grid. Model validation was performed using two complementary approaches: ten-fold cross-validation assessed predictive accuracy by repeatedly partitioning the dataset into ten subsets, training the model on nine subsets, and evaluating performance on the remaining subset, thus providing mean and standard error estimates for each predictive metric across all folds. Additionally, out-of-bag validation was used to obtain an internal estimate of model accuracy, based on predictions made for each observation using only the subset of trees that excluded it during training. This approach provided an unbiased estimate of the coefficient of determination without requiring a separate validation set. We chose random cross rather than block cross-validation (Roberts et al. 2017) because our goal was to model species richness along the most continuous spatial and temporal gradients possible, and enforcing blocks on coordinates would have prevented the algorithm from learning those gradients effectively.

Variable importance scores for the final models were estimated using permutation and adjusted for the R² of each model. For an *i*-th predictor, the formula is:

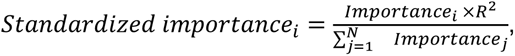

Where *i* and *j* index individual predictors, Importance*_i_* is the permutation importance value of the i-th predictor, N is the total number of predictors, and R² is the coefficient of determination of the model. This approach ensures that the importance values are scaled relative to the model’s explanatory power, providing more intuitive comparisons across models. We also quantified variable interactions for each model using hstats version 1.2.1 (Mayer 2024), computing pairwise interaction strengths (H² statistics) and overall interaction contributions. All models were adjusted and implemented using a tidymodels workflow (Kuhn and Wickham 2020), utilizing the R packages tidymodels version 1.2.0 (Kuhn et al. 2022), ranger version 0.16.0 (Wright and Ziegler 2017), and vip version 0.4.1 (Greenwell and Boehmke 2020).

We used partial dependence plots (PDPs) to visualize how key predictors affect bird species richness in our random forest models using the R package hstats (Mayer 2024). PDPs estimate the marginal effect of a predictor by varying it across its range. For each value *x* of a focal predictor *i*, predictions are made across the entire dataset by substituting *x* for the original observed values of predictor *i* in every observation, while keeping all other predictors at their original observed values. The resulting predictions are then averaged, allowing the visualization of the influence of the focal predictor. We generated PDPs for each model, showing how the effects of the predictors change across different time spans. With this approach, we show the direct, time-span-dependent effects of each environmental variable on species richness.

### Varying sampling effort

Species richness tends to vary with varying sampling effort during a survey, which may affect our conclusions. We do not expect this to affect analyses using the JPSP dataset, as the sampling effort was standardized. We have no information about the sampling effort for the SPARSE dataset, and thus make the assumption that it is captured by the duration and area of a survey, but this also implies that results based solely on the SPARSE data should be interpreted with some caution.

The Atlases dataset has a standard survey protocol, but the effort (i.e., number of observers per cell and number of their visits) can still vary. To assess model sensitivity to this variation, we incorporated sampling effort as a predictor for this dataset using the Frescalo algorithm (Hill 2012). This method allows the estimation of recorder effort based on species’ local frequency patterns in spatially contiguous neighborhoods, a condition met only by the Atlases dataset. For this dataset, effort estimates were calculated separately for each sampling period and averaged across the grouped combinations. Because incorporating the effort as a predictor did not alter the patterns observed in our Atlases model, we present the result only as a value importance plot in Supplementary Fig. 5.

### Predictions from random forests over different time spans

To examine how different time spans affect predictions of a macroecological model of species richness, we predicted bird species richness across Czechia using the JPSP points dataset for three time spans: 1-year, 5-year, and 15-year sampling windows. These time spans were selected because they approximate common temporal spans used in biodiversity monitoring schemes (e.g., annual, short–term, and long–term programmes) and are well represented in our datasets (Fig. 1). We selected this JPSP points-based model because the spatial and temporal distribution of the sampling points was more spatially homogeneous than in the other datasets. Predictions were generated over a set of artificial 12.6 km² circular points (equivalent to ∼2 km radius), spaced 10 km apart between cell centers, and covering the entire country. We chose the values for the year 2010 for other covariates (land use and climatic predictors) because this year was the median sampling year in the JPSP dataset (sampling period: 1985–2020). For the 1-year time span, we used only data from 2010 for all predictor variables. For the 5-year and 15-year time spans, we used variable values for 2010 and from 2008-2012 and 2003-2017, respectively, extracting or averaging predictor values across the corresponding years around the central reference year. For each time span, we then identified richness hotspots and coldspots as the 10% of grid cells with, respectively, the highest and lowest predicted species richness values.

## Results

### Model performance

All models identified time span, and area as the most important variables explaining variation in bird species richness, although their relative ranking varied across datasets (Fig. 2a). On average, these three variables accounted for 20-60% of the total importance, independently of the dataset used. Among climatic variables, precipitation was generally the most important, except in the Atlases model, where minimum temperature dominated, while among land-cover variables, water bodies showed the highest importance across nearly all models. Pairwise interaction estimates (H²) indicated that time span and area showed strong interactions with other predictors, whereas their direct interaction with each other was comparatively weak (Fig. 2b, Supplementary Fig. 6). Model performance was high (i.e., cross-validation and out-of-bag), with R² values ranging from 0.44 (Atlases) to 0.95 (All datasets) (Supplementary Table 2, Supplementary Fig. 4).

**Figure 2.**
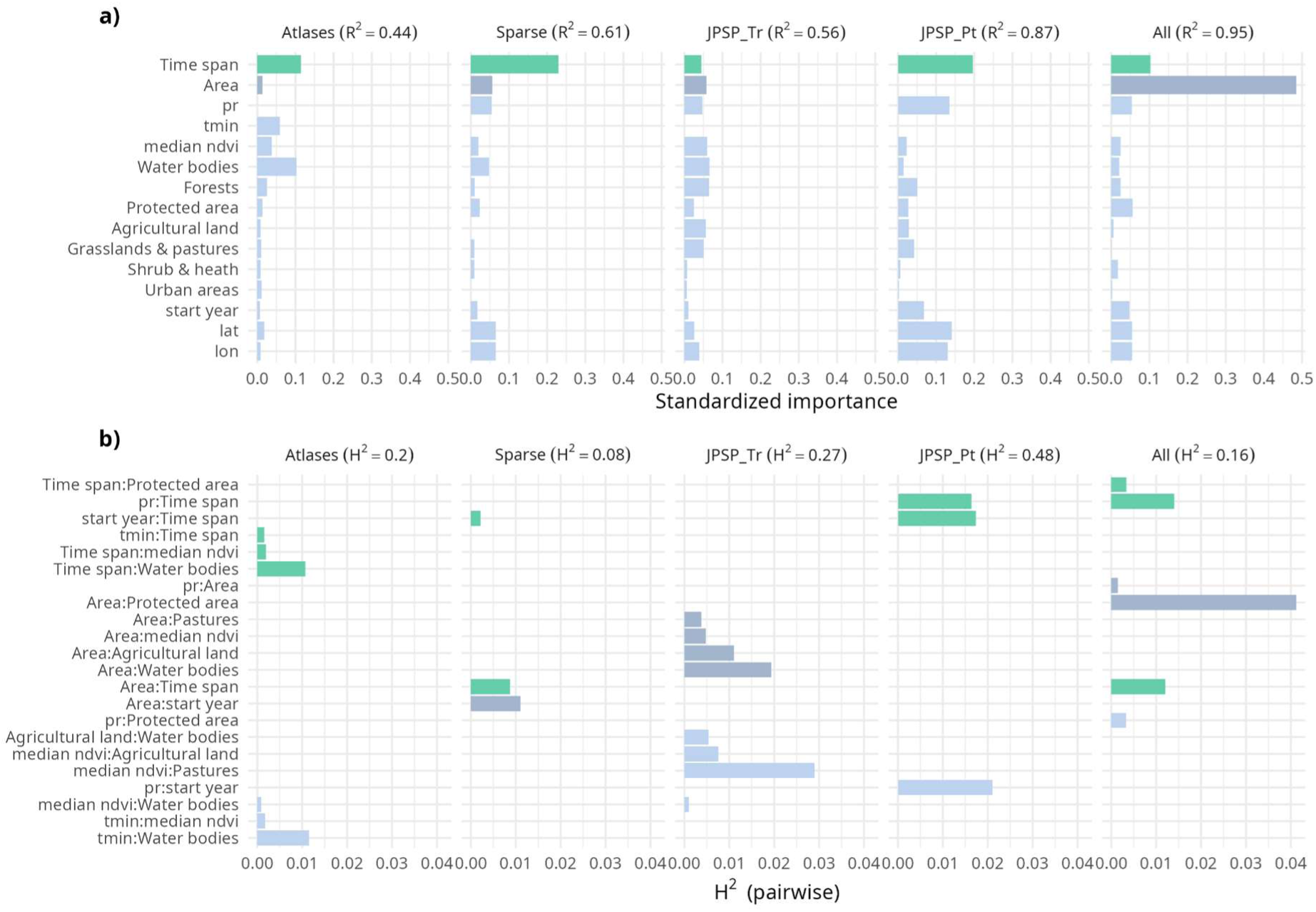
a) Relative importance of predictors in random forest models explaining variation in species richness across Czechia. Bars represent the relative importance of each variable (see methods). (b) Pairwise contributions to explained variance (H^2^) for the most influential two-way interactions in each model (see supplementary Fig. 6 for all interactions). The panels correspond to random forest models fitted separately for each dataset (i.e., Atlases, SPARSE, and JPSP), and the last panel represents the model that uses all the data together. Green bars correspond to the Time span, grey bars represent the Area, and light blue bars represent all remaining variables/interaction terms.

### Species time-area relationship (STAR)

Across all models, both area and time span showed positive effects on species richness, and their interaction revealed a consistent STAR pattern: larger areas and longer time spans were associated with higher species richness. The curves indicate that the combined effects of space and time are multiplicative rather than additive, with longer time spans increasing richness across all area sizes, and short time spans showing the steepest species–area gains at small spatial grains (Fig. 3).

**Figure 3.**
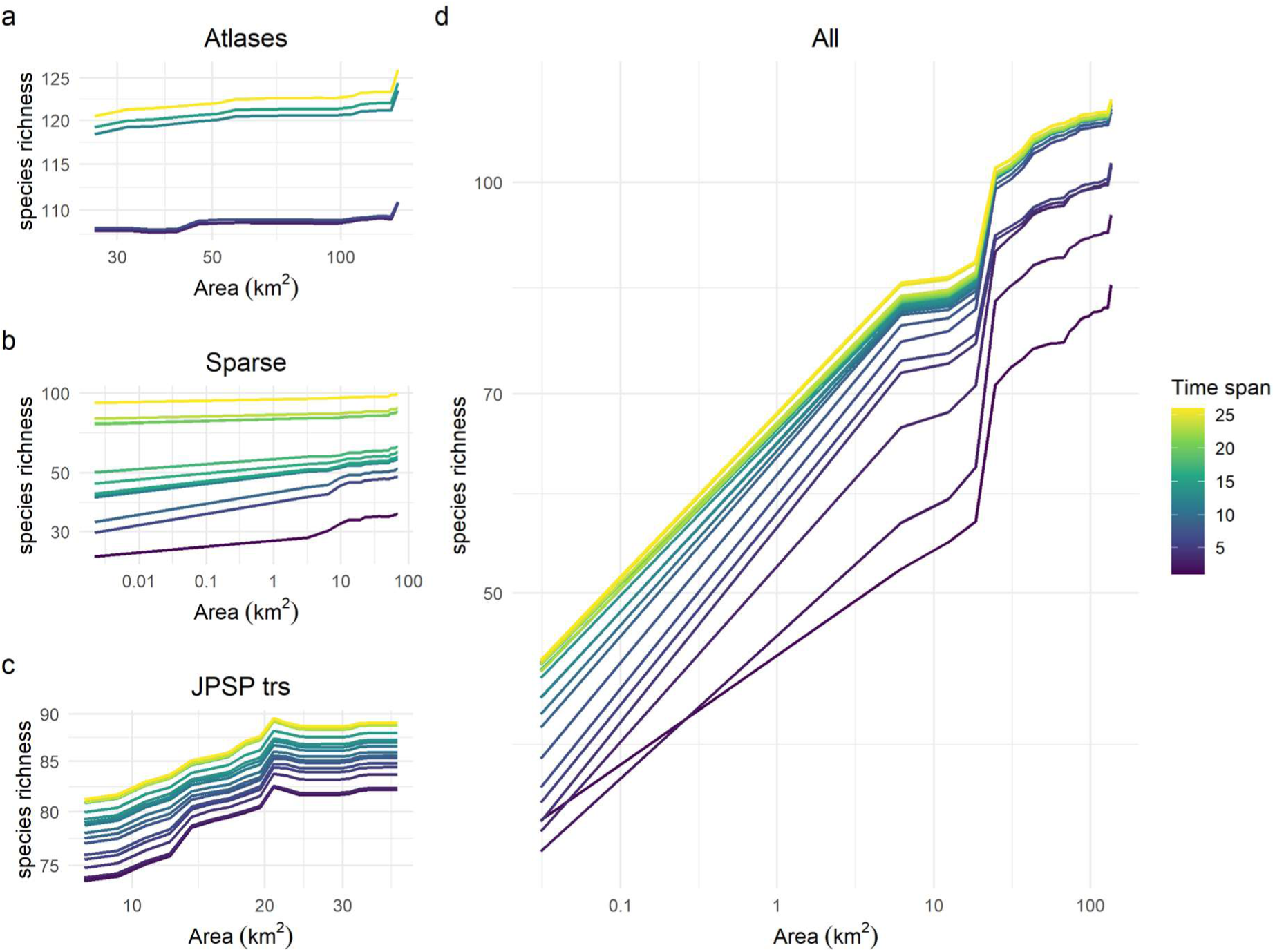
Partial dependence plots of the relationship between species richness and area across different time spans. All variables were plotted on a logarithmic scale, but untransformed values were shown as labels to improve interpretability.

### Drivers of species richness

Our models revealed that precipitation and water bodies are the key environmental predictors of bird species richness, both in terms of variable importance and interactions with time and area. In the case of mean annual precipitation, species richness decreased, with steeper slopes around 600 mm y^-1^ (Fig. 4). Furthermore, the presence of water bodies and their size positively influenced species richness in all of our models, with steeper slopes in areas with smaller water bodies, which tended to stabilize at around 0.7 km² in most models (Fig. 5). The interaction effects of time span with these predictors on species richness was positive, with curves of longer time spans consistently related to higher species richness than shorter time spans, regardless of the increase in area of water bodies or precipitation amount (Figs. 4 and 5).

**Figure 4.**
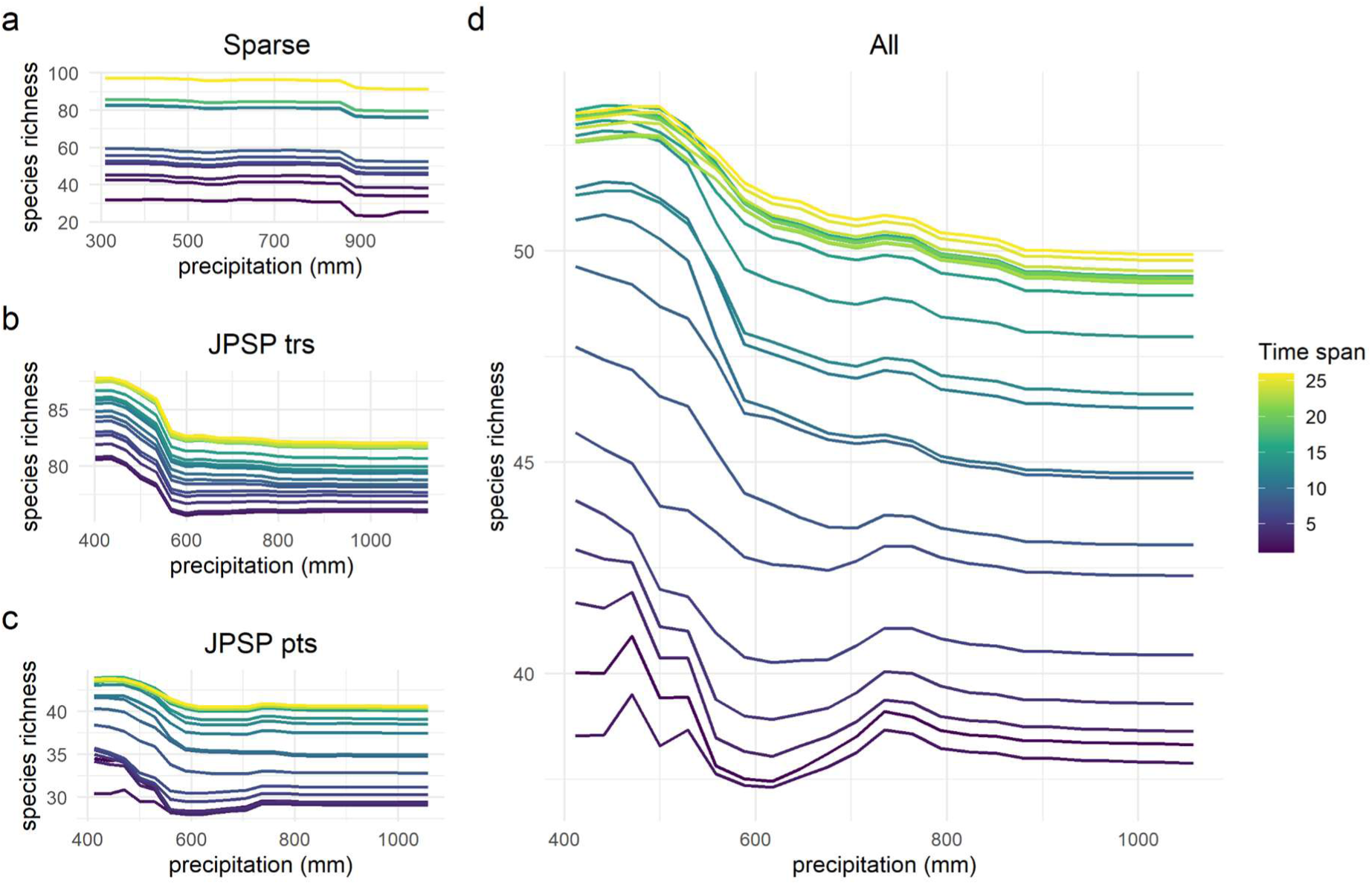
Partial dependence plots of the relationship between species richness and precipitation (i.e., the most important climatic predictor for these four models, see figure 2) across different time spans. Each panel corresponds to a different random forest model fitted on a different dataset. In the case of the plot titled “All”, this corresponds to the model that includes all data sets (Sparse, JPSP, and Atlases).

**Figure 5.**
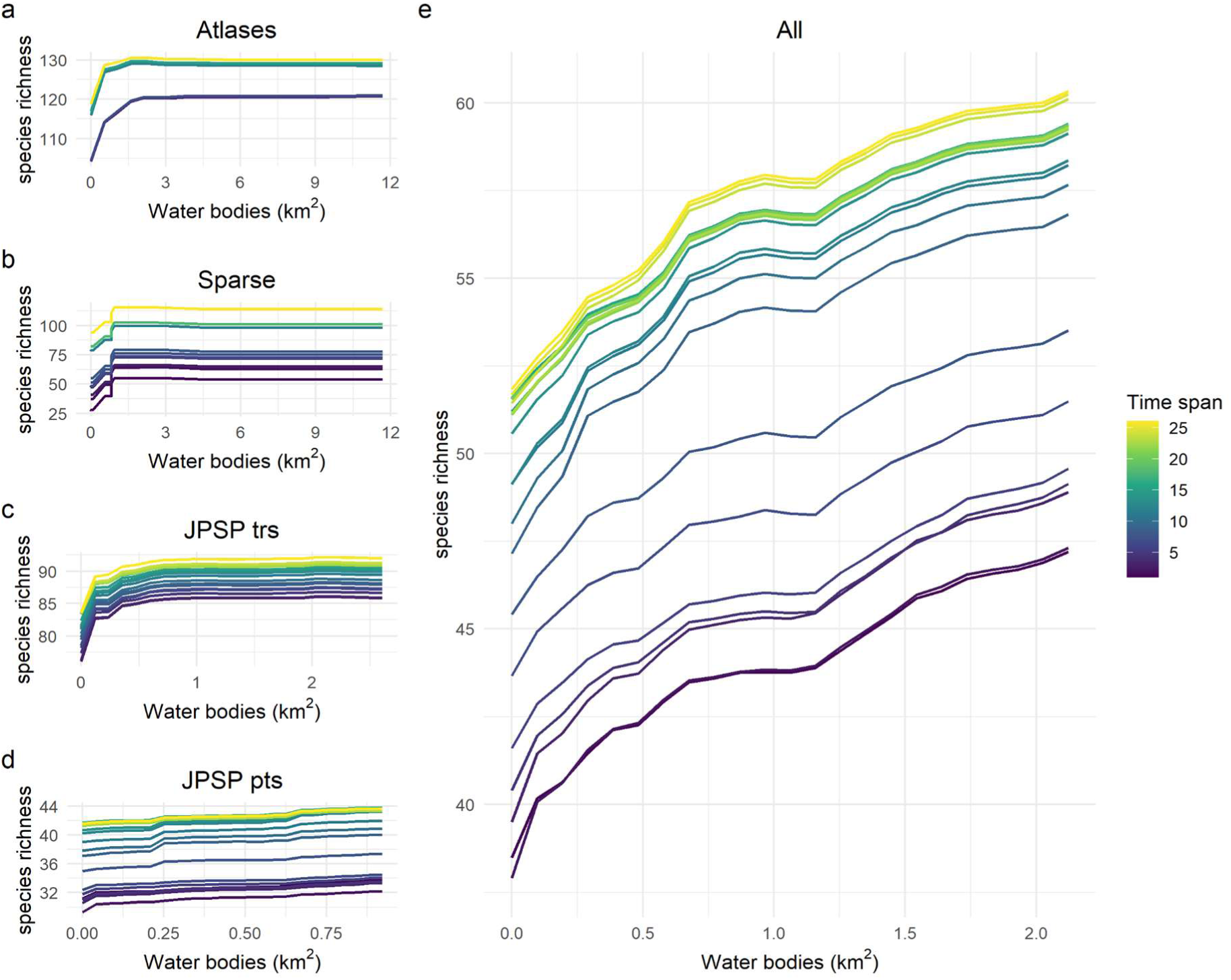
Partial dependence plots of the relationship between species richness and the size of water bodies within the sampling unit across different time spans (in years; a-d). In e, this corresponds to the Random Forest model that includes all data sets (Sparse, JPSP, and Atlases).

### Predicted hotspots of bird diversity

The predictions made for three time spans showed notable variations in modeled bird species richness (Fig. 6). On one hand, the predicted species richness for one-year and five-year time spans was relatively similar, but the hotspots of species richness (i.e., the 10% of the points with the highest species richness) shifted to different regions when different time spans were used. Specifically, these hotspots shifted from being concentrated near the borders of Czechia when estimated over shorter time spans, to being more concentrated in central Czechia when estimated over longer time spans. Conversely, coldspots (i.e., the 10% of the points with the lowest species richness) tended to be clustered in the center and south of Czechia, and gradually shifted towards border regions over longer time spans.

**Figure 6.**
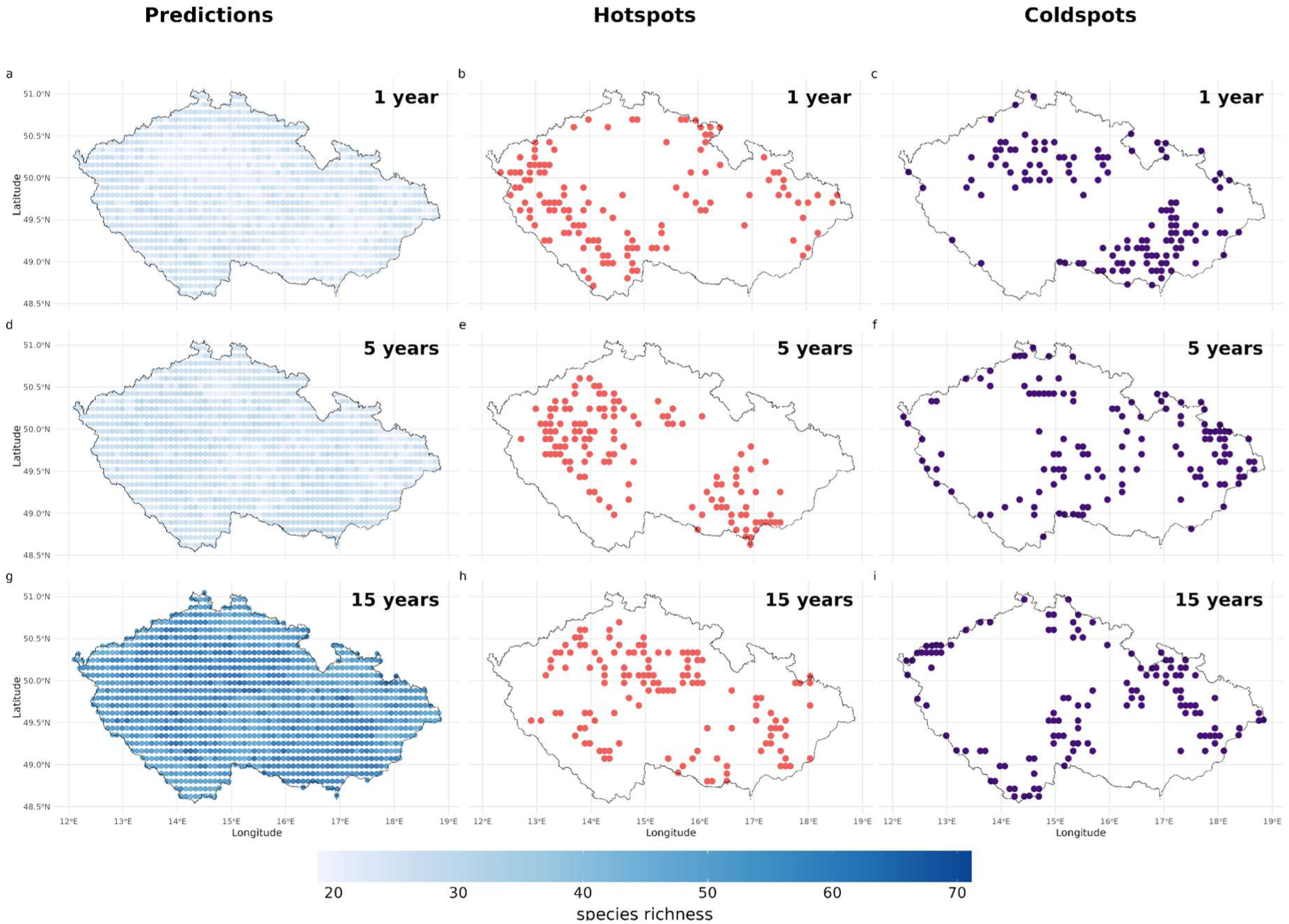
Predicted bird species richness in Czechia, based on the Random Forest model of JPSP points using three different time spans: 1 year (A, B, C), 5 years (D, F, G), and 15 years (G, H, I). Orange circles represent the hotspots of species richness (i.e., 10% of the points with the highest bird species richness) and purple circles represent the coldspots of species richness (i.e., 10% of the points with the lowest bird species richness).

The inclusion of the sampling effort in the Atlases model did not substantially change our results, and the time span consistently remained the most important variable (supp. Fig. 5), supporting the robustness of our conclusions.

## Discussion

We found that that time span was consistently ranked among the most important predictors of bird species richness. Furthermore, in datasets with the widest range of time spans and area, it also had important interactions with area and key environmental predictors such as precipitation and water bodies. Although this may reflect the well-known link between sampling effort (duration in minutes or hours) and observed diversity (Gotelli & Colwell 2001), our results expand this concept to much longer time frames from years to decades, revealing significant annual turnover in bird assemblages. Furthermore, the effect of time span on other drivers of species richness highlights the need to explicitly account for time span when evaluating and predicting species richness patterns.

### Joint effects of time span and area

We found a strong positive association between area and species richness (Fig. 3), consistent with the classic species-area relationship (Preston 1960, Adler and Lauenroth 2003, Adler et al. 2005), where larger areas support higher species diversity due to the increased availability of resources and habitat heterogeneity. In addition, for the datasets with the widest range of values of both area and time span (i.e., the SPARSE dataset and all datasets combined), area also had a pairwise interaction with time span (Fig. 2b, Fig. 3b, d). This combined pattern (Fig. 3) shows that temporal grain systematically increases species richness across the full range of spatial scales and regulates the steepness of the STAR at short time spans. Previous studies (Rahbek 2005, Field et al. 2009, Belmaker and Jetz 2011, Keil and Chase 2019) have established that spatial scale is critically important in biodiversity, and we confirm that temporal span acts as a complementary axis that modulates the influence of area on species richness. These findings mirror those found by Adler et al. (2005), where grassland plant communities exhibited steeper SAR slopes in short-term surveys compared to long-term surveys.

### Effect of environment and its interaction with the time span

Precipitation and area of water bodies were the best environmental predictors of bird species richness across most of our models, in line with previous studies on the drivers of bird species richness in Czechia (Storch et al. 2003, Dvořáková et al. 2023). Precipitation showed a slightly negative relationship with species richness, likely due to colder conditions at higher, wetter sites, a pattern also observed in the country for plant species richness (Pyšek et al. 2002). Conversely, the presence of water bodies had a positive effect on richness, implying that these habitats promote specialists and contribute to greater avian biodiversity across both urban and natural ecosystems (Chamberlain et al. 2007, Xie et al. 2022, Cerda-Peña and Rau 2023, Dvořáková et al. 2023, Aubrechtová et al. 2024). Even small water bodies significantly increased species richness, consistent with saturation in the use of vegetated littoral zones rather than the entire aquatic surface (Pšeničková and Horák 2022, Gábor et al. 2023).

Importantly, in datasets with the widest range of time span values (i.e., SPARSE, all datasets combined), these predictors had an interaction effect with time span, (i.e., time span steepened their relationships with species richness) (Figs. 4b, 4d, 5a), and this was particularly obvious in the effect of precipitation. Specifically, longer time spans were consistently associated with higher species richness than shorter time spans, regardless of precipitation amount or water body area (Figs. 4, 5). These interactive effects between the time span and both precipitation and water bodies on species accumulation likely arise because longer time spans can capture the responses of species to climatic and habitat fluctuations, as well as episodic or rare occurrences over time, that short time spans tend to miss.

### Shifting predicted hotspots

We found that the predicted spatial patterns of bird species richness varied significantly when different time spans were used. As expected, average species richness increased with the length of the time span. While we did not detect marked differences in species richness between one-year and five-year time spans (Fig. 6a, d), we found a striking increase in species richness at time spans of 15 years (Fig. 6e). In addition, the spatial patterns of species richness across Czechia shifted as the timespan increased, suggesting that time could affect diversity in different ways than area.

Our predictions using a 1-year time span showed that hotspots were concentrated along the west and east borders of Czechia, especially in the southwest, where wetlands and ponds are abundant and provide heterogeneous habitats that can contribute to enhancing species richness. Using a 5-year time span, new hotspots emerged in the southeast, particularly in lowland wet forests, which are known as some of the most biologically diverse regions in Czechia, especially for insects (Miklín et al. 2017). This shift in the location of hotspots may suggest that species richness in water bodies accumulates rapidly, whereas forest communities tend to build up more gradually and may also be harder to detect than those using open water. In contrast, 15-year time span predictions (Fig. 6h) showed hotspots shifting to central lowland agricultural areas, which previously had been classified as coldspots when using predictions with 1-year timespans (Fig. 6c). This may indicate the presence of species that are rare, specialists, or more difficult to detect, possibly due to the higher fragmentation and heterogeneity of the landscape resulting from human impacts (Austen et al. 2001, Devictor et al. 2008, Harrison et al. 2019). The intensively farmed agricultural lands associated with these locations may indicate a reduced capacity for these areas to support diverse bird communities in the short term (Dvořáková et al. 2022). Yet, after 15 years, these same regions become bird diversity hotspots, as also documented previously in Europe (Anderle et al. 2023, Edo et al. 2023). This pattern could also be explained by the more favorable climatic conditions and landscape connectivity of lowland sites, which may facilitate species accumulation over time compared to the harsher, more isolated conditions of highland forests (Fig. 6i).

### Implications for monitoring and conservation

Our findings have practical implications for biodiversity research and conservation. When comparing diversity across sites, it is crucial to standardize the sampling duration, just as it is frequently done for spatial grain; when this is not feasible, we suggest recording the time span and including it as a covariate in statistical models to control for temporal effects and improve ecological inferences.

We have also demonstrated that longer sampling periods not only detect higher species richness but also different biodiversity hotspots compared to shorter sampling periods, suggesting that short-term surveys may underestimate diversity and overlook critical conservation areas. This is particularly crucial in the context of climate change, which is rapidly altering habitats, migration patterns, and species distributions (Huntley et al. 2006, Chen et al. 2011, Bateman et al. 2016). Effective conservation strategies should therefore prioritize extensive, long-term biodiversity surveys and explicitly consider monitoring duration when identifying priority areas, detecting vulnerable or elusive species, and adapting management actions to ongoing environmental changes.

## Conclusion

We demonstrated that time is not just a background feature of sampling; it is a fundamental axis of biodiversity. Just as spatial scale has long been recognized as central to ecological theory (Wiens 2000, Sandel 2015), temporal scale emerges as a key dimension that modulates both the magnitude and the spatial configuration of species richness and its hotspots. Short-term surveys can mask long-term diversity trends and prioritize conservation areas that may not be high in species richness. Conversely, long-term observations reveal ecological processes that determine the accumulation of diversity over time. Thus, biodiversity can only be studied comprehensively at multiple temporal grains simultaneously, i.e., by using the species-time relationship, or even better, the species-time-area relationship. Integrating time span as a fundamental dimension alongside spatial grain enhances the comparability of biodiversity patterns across studies, strengthens macroecological inference, and supports the development of conservation strategies in the context of ongoing global change.

## Data availability

The atlases data are not yet freely available and were provided for this study by K. Šťastný and V. Bejček. The JPSP data were provided for this study by J. Reif, Z. Vermouzek, Petr Voříšek, K. Šťastný, V. Bejček, and J. Janda.

## Supplementary information

**Table S1.** Summary of the final predictors used in each Random Forest model after checking for collinearity (see Methods section). The “x” indicates the predictors included in each final model. All models included as predictors time span, start year, area, latitude, and longitude. *Land principally occupied by agriculture with significant areas of natural vegetation.

| Predictor | Atlases | Sparse | JPSP_pts | JPSP_trs | All |
| --- | --- | --- | --- | --- | --- |
| Precipitation |  | x | x | x | x |
| Temperature min | x |  |  |  |  |
| Temperature max |  |  |  |  |  |
| Mean Temperature |  |  |  |  |  |
| Median ndvi | x | x | x | x | x |
| Broad-leaved forest | x | x | x | x | x |
| Complex cultivation patterns | x |  | x | x | x |
| Construction sites | x | x | x | x | x |
| Coniferous forest | x |  | x |  |  |
| Continuous urban fabric | x | x | x | x |  |
| Discontinuous urban fabric | x |  |  | x |  |
| Green urban areas |  | x | x | x | x |
| Agricultural land* | x |  | x | x |  |
| Mixed forest | x |  | x | x |  |
| Moors and heathland | x | x | x | x | x |
| Natural grasslands | x |  | x | x | x |
| Pastures | x | x | x | x |  |
| Transitional woodland shrub | x | x | x | x | x |
| Water bodies | x | x | x | x | x |
| Water courses | x | x | x | x | x |
| Protected area | x | x | x | x | x |

**Table S2.** Summary of Random Forest regression performance metrics for bird species richness in Czechia, assessed using two validation methods: 10-fold cross-validation (CV) and out-of-bag (OOB) estimates. For cross-validation, the coefficient of determination (R²), root mean squared error (RMSE), and mean absolute error (MAE) were calculated using the fit_resamples() function in the Tidymodels R package, reporting the mean and standard error across test folds. OOB metrics—OOB R² and OOB mean squared error (OOB MSE)—were obtained from the ranger engine, reflecting model performance based on predictions for observations excluded from each tree during training.

| Model | $R^2$ | RMSE | MAE | OOB $R^2$ | OOB MSE |
| --- | --- | --- | --- | --- | --- |
| All | 0.928 ± 0.004 | 7.581 ± 0.192 | 4.741 ± 0.076 | 0.931 | 55.25 |
| JPSP Points | 0.861 ± 0.004 | 4.730 ± 0.061 | 3.376 ± 0.052 | 0.867 | 21.32 |
| JPSP Transects | 0.557 ± 0.038 | 13.791 ± 0.901 | 9.979 ± 0.718 | 0.514 | 190.14 |
| Atlases | 0.503 ± 0.321 | 14.142 ± 0.487 | 11.185 ± 0.321 | 0.503 | 198.29 |
| SPARSE | 0.685 ± 0.046 | 16.738 ± 2.861 | 10.59 ± 1.561 | 0.586 | 341.42 |

**Supplementary Figure 1.**
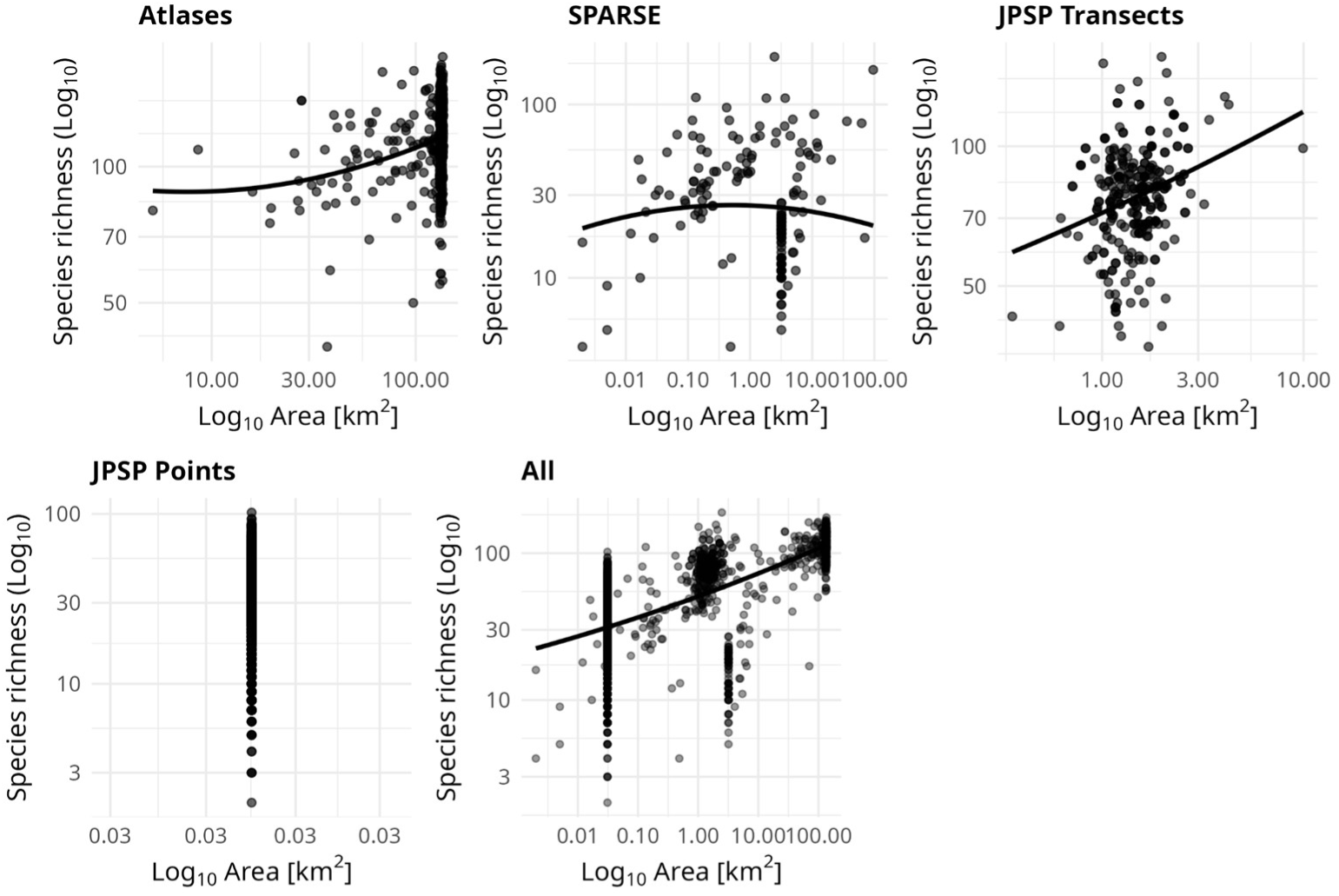
Species richness-area relationship for each data set used in this study.

**Supplementary Figure 2.**
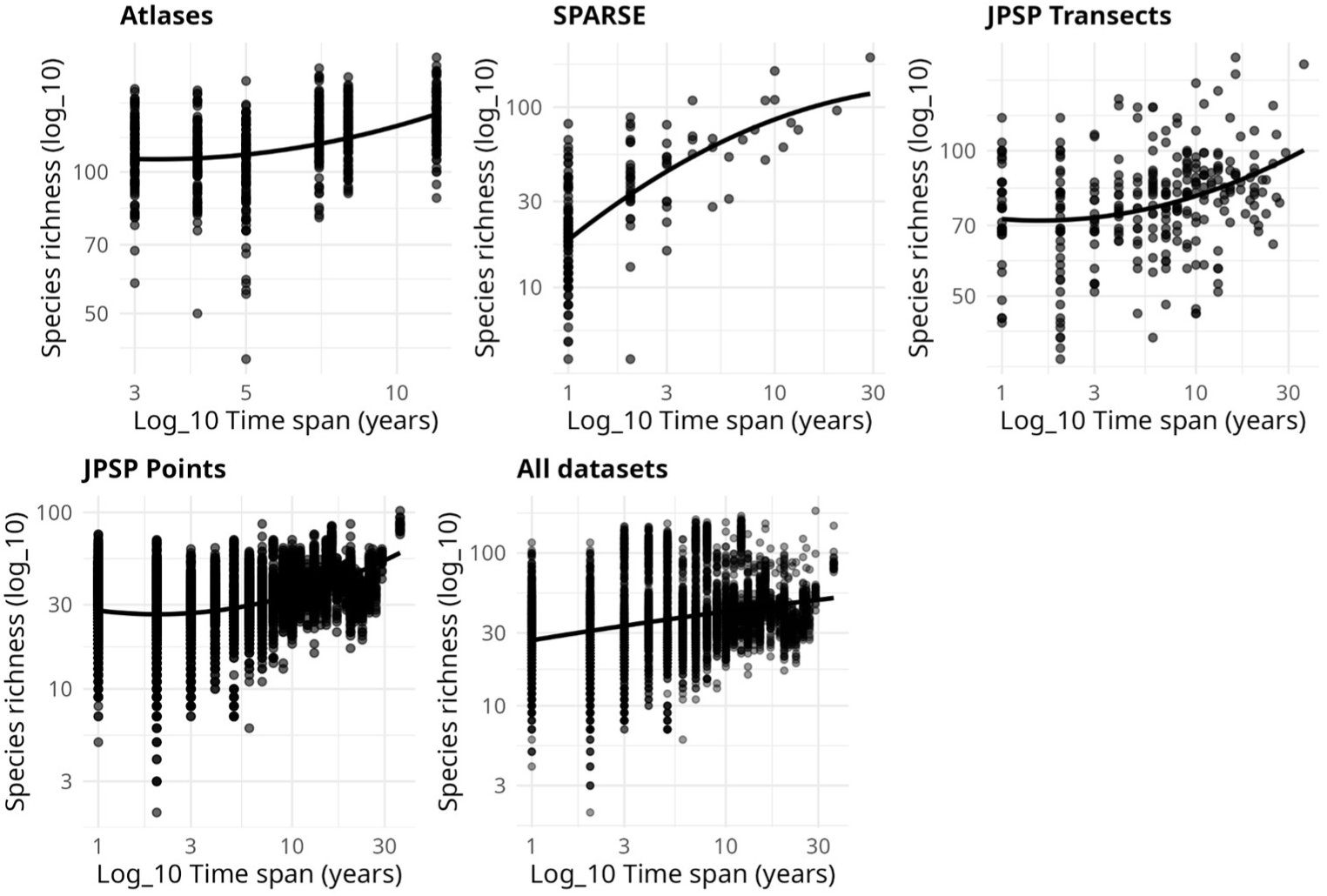
Species richness-time relationship for each data set used in this study.

**Supplementary Figure 3.**
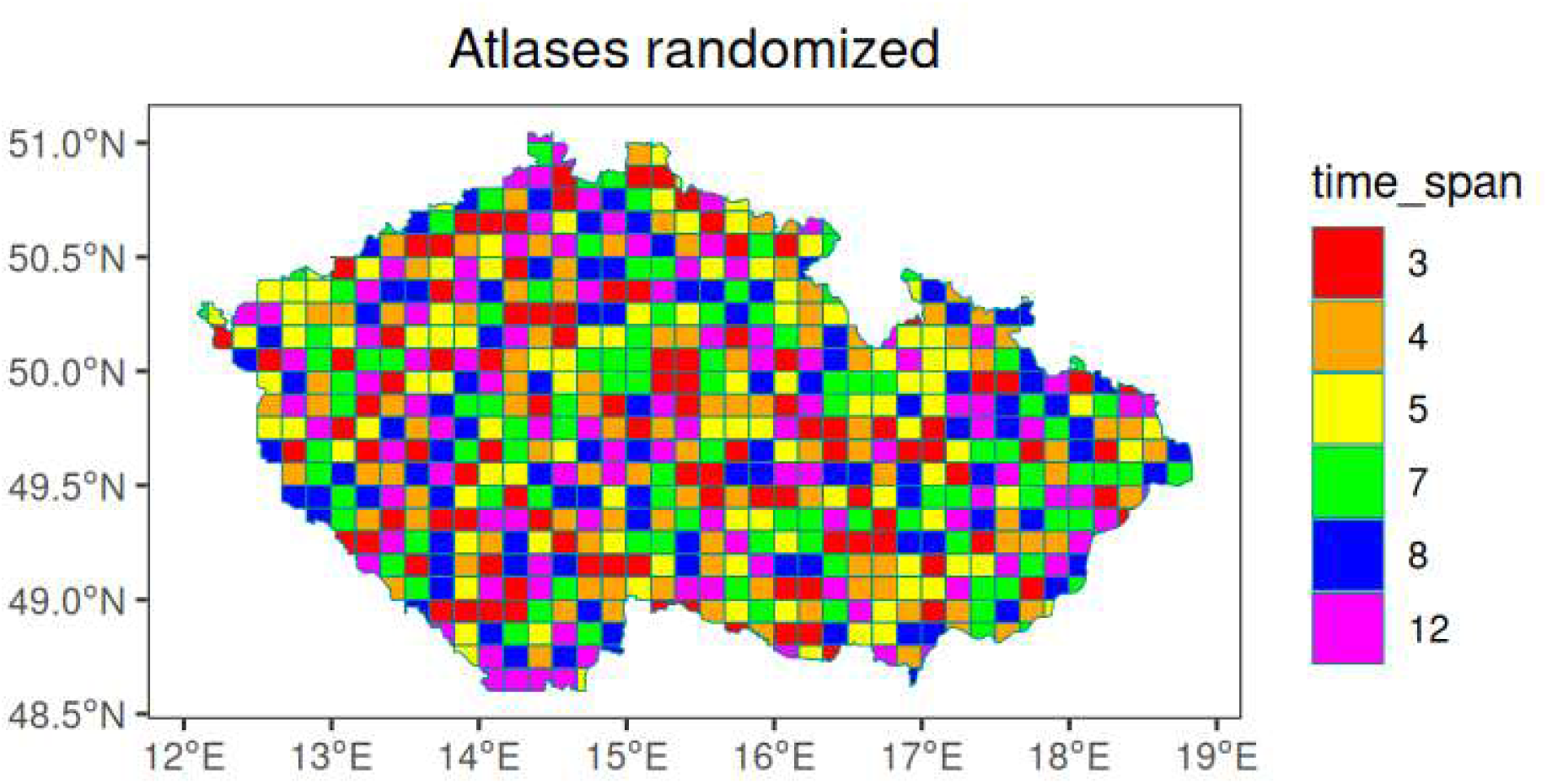
The atlases dataset mapped to show the different time-spans of the six groups of cells: 3= Atlas II (2001-2003), 4= Atlas III (2014-2017), 5= Atlas I (1985-1989), 7= Atlas II+ Atlas III, 8= Atlas I + Atlas II, 12= Atlas I + Atlas II+ Atlas III.

**Supplementary Figure 4.**
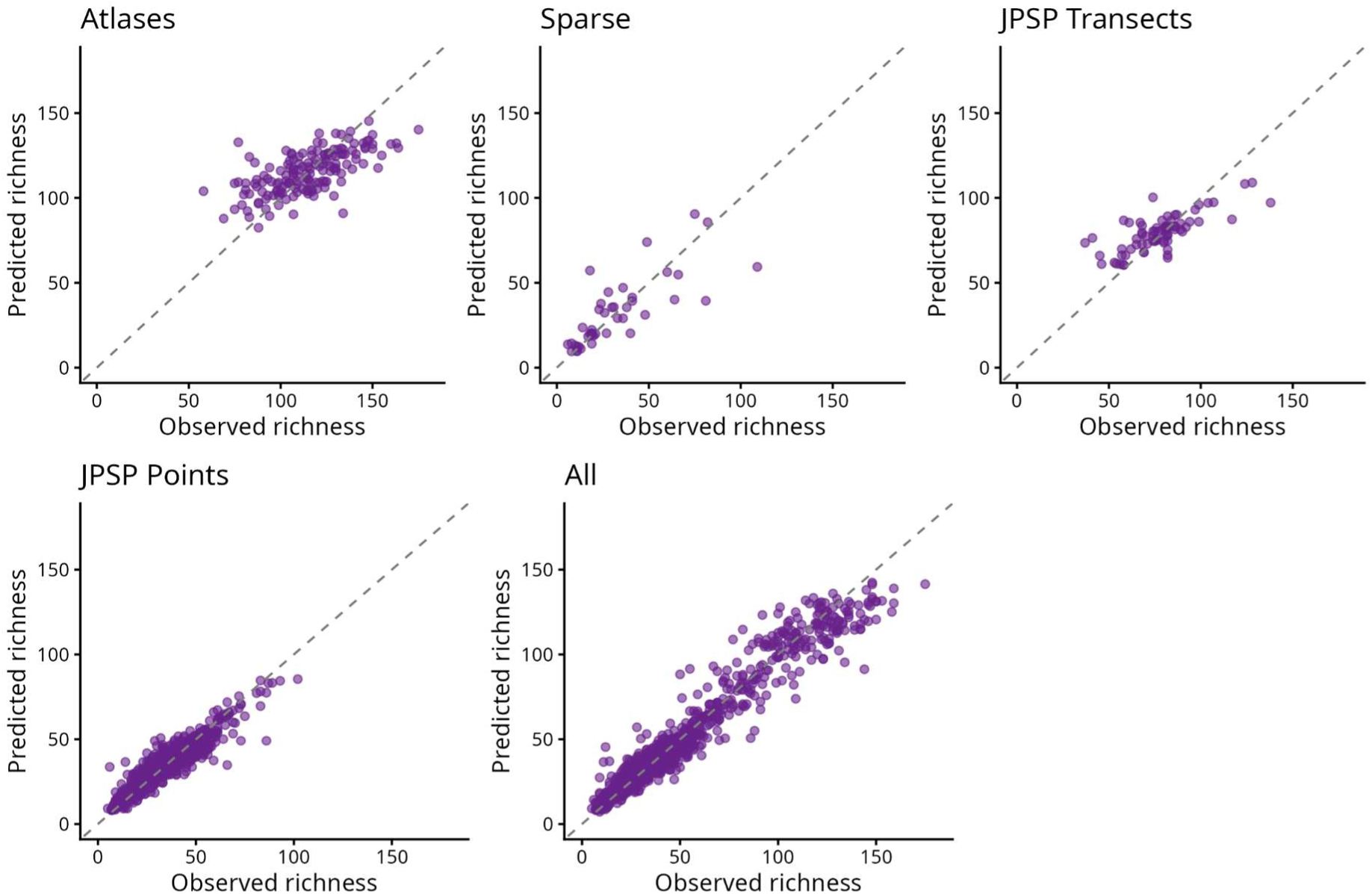
Observed versus predicted species richness values for the test set of all the Random Forest models. Each point represents a test observation; the dashed diagonal line indicates the perfect 1:1 match between observed and predicted richness.

**Supplementary Figure 5.**
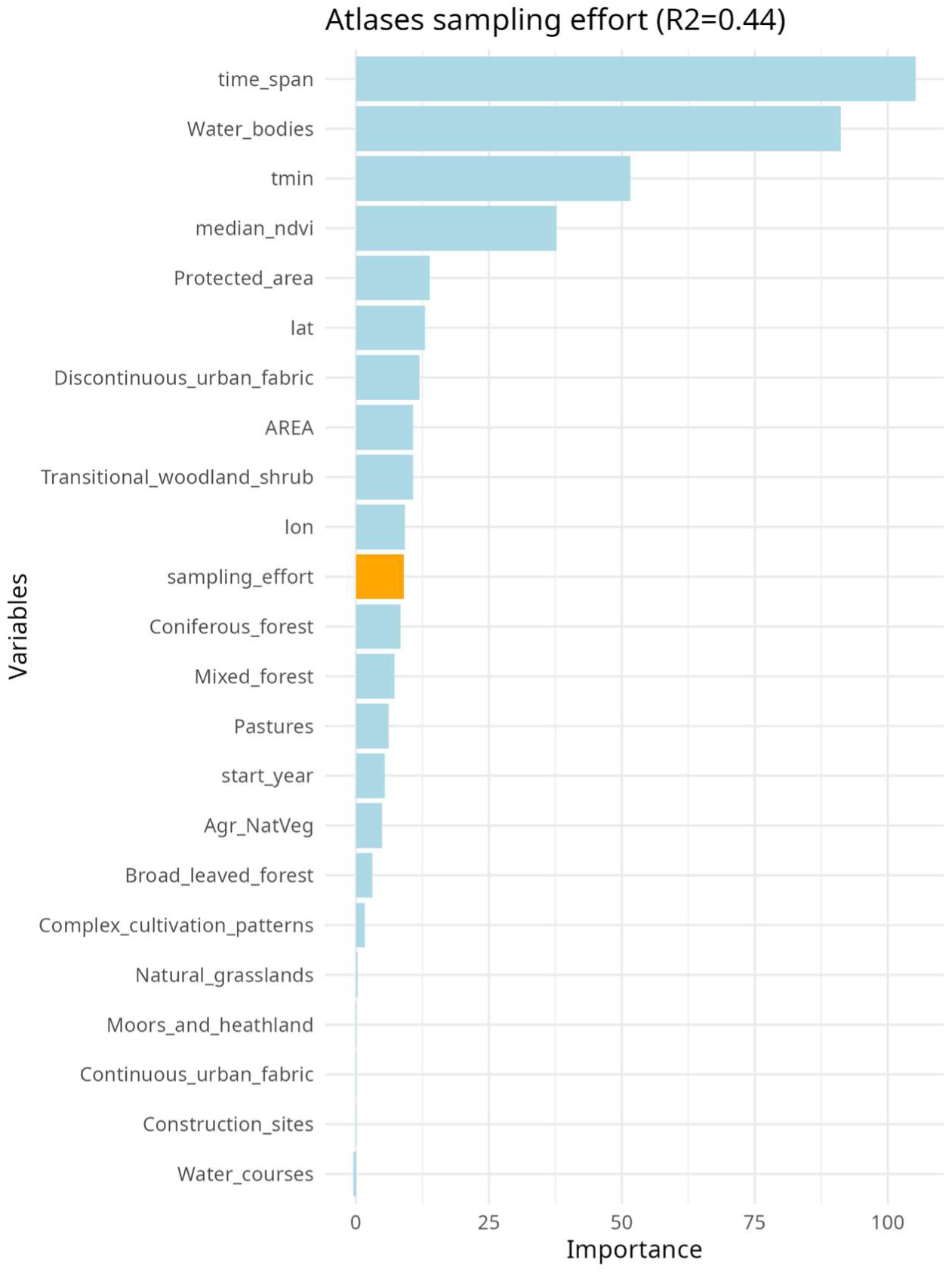
Variable importance plot incorporating the sampling effort into our Atlases model. The sampling effort (orange bar) was calculated using the Frescalo algorithm.

**Supplementary Figure 6.**
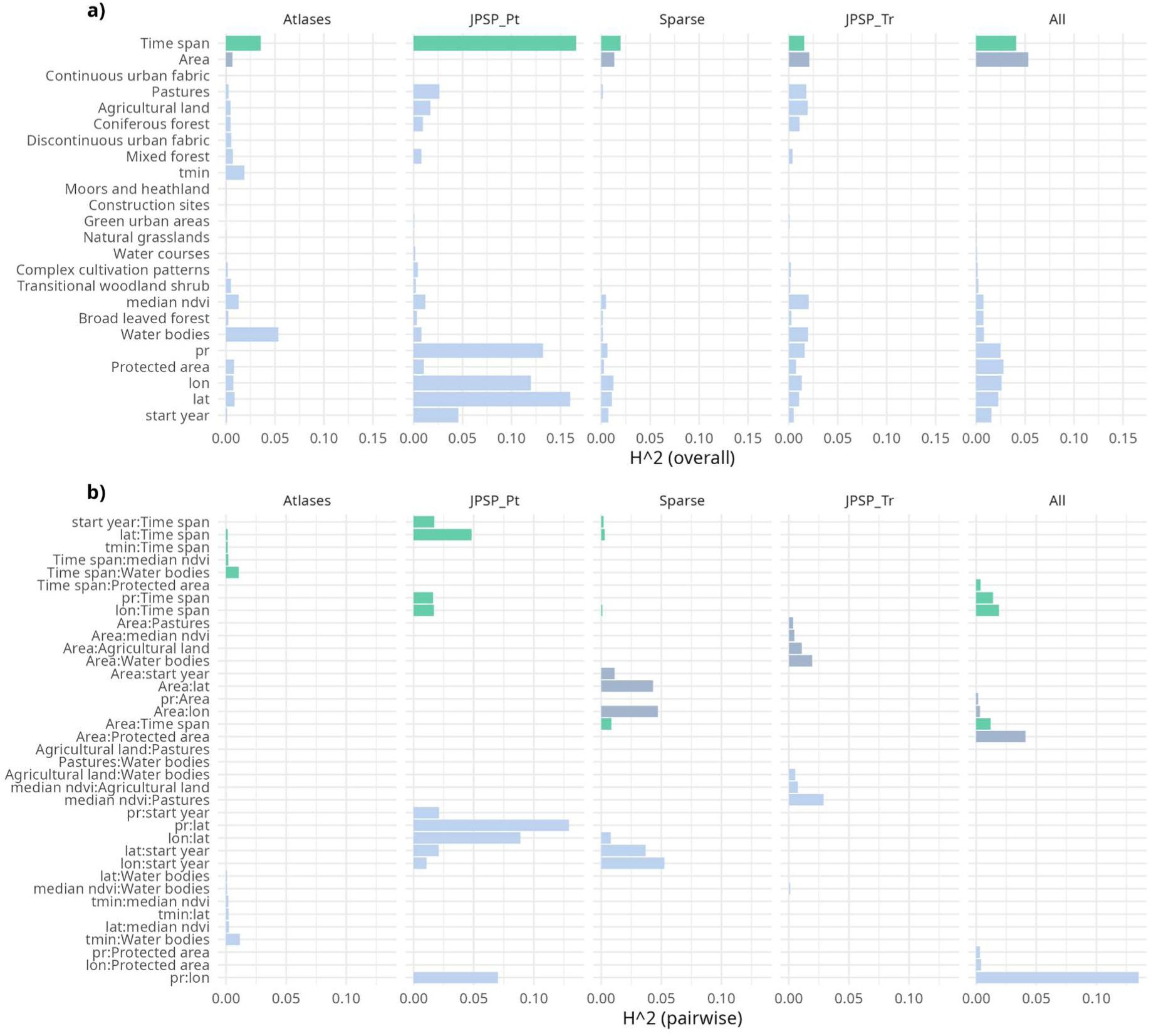
Overall and pairwise interactions calculated using HSTAT. Panel a) reports the proportion of prediction variability explained by interactions on a predictor. Panel b) reports the proportion of joint effect variability of features coming from their pairwise interaction.

## References

1. Adler, P. B. and Lauenroth, W. K. 2003. The power of time: spatiotemporal scaling of species diversity. - Ecol. Lett. 6: 749–756.

2. Anderle, M., Brambilla, M., Hilpold, A., Matabishi, J. G., Paniccia, C., Rocchini, D., Rossin, J., Tasser, E., Torresani, M., Tappeiner, U. and Seeber, J. 2023. Habitat heterogeneity promotes bird diversity in agricultural landscapes: Insights from remote sensing data. - Basic Appl. Ecol. 70: 38–49.

3. Aubrechtová, E., Bydžovská, T. and Horák, J. 2024. Blue-green infrastructure and biodiversity: Urbanization and forestation have an important influence on bird diversity in water habitats. - Urban For. Urban Green. 91: 128151.

4. Austen, M. J. W., Francis, C. M., Burke, D. M. and Bradstreet, M. S. W. 2001. Landscape Context and Fragmentation Effects on Forest Birds in Southern Ontario. - Condor Ornithol. Appl. 103: 701–714.

5. Bateman, B. L., Pidgeon, A. M., Radeloff, V. C., VanDerWal, J., Thogmartin, W. E., Vavrus, S. J. and Heglund, P. J. 2016. The pace of past climate change vs. potential bird distributions and land use in the United States. - Glob. Change Biol. 22: 1130–1144.

6. Belmaker, J. and Jetz, W. 2011. Cross-scale variation in species richness–environment associations. - Glob. Ecol. Biogeogr. 20: 464–474.

7. Breiman, L. 2001. Random Forests. - Mach. Learn. 45: 5–32.

8. Catford, J. A., Wilson, J. R. U., Pyšek, P., Hulme, P. E. and Duncan, R. P. 2022. Addressing context dependence in ecology. - Trends Ecol. Evol. 37: 158–170.

9. Cerda-Peña, C. and Rau, J. R. 2023. The importance of wetland habitat area for waterbird species-richness. - Ibis 165: 739–752.

10. Chamberlain, D. E., Gough, S., Vaughan, H., Vickery, J.A. and and Appleton, G. F. 2007. Determinants of bird species richness in public green spaces. - Bird Study 54: 87–97.

11. Chen, I.-C., Hill, J. K., Ohlemüller, R., Roy, D. B. and Thomas, C. D. 2011. Rapid Range Shifts of Species Associated with High Levels of Climate Warming. - Science 333: 1024–1026.

12. Colwell, R. K. and Coddington, J. A. 1994. Estimating terrestrial biodiversity through extrapolation. - Philos. Trans. R. Soc. Lond. B. Biol. Sci. 345: 101–118.

13. Craven, D., van der Sande, M. T., Meyer, C., Gerstner, K., Bennett, J. M., Giling, D. P., Hines, J., Phillips, H. R. P., May, F., Bannar-Martin, K. H., Chase, J. M. and Keil, P. 2020. A cross-scale assessment of productivity–diversity relationships. - Glob. Ecol. Biogeogr. 29: 1940–1955.

14. Devictor, V., Julliard, R. and Jiguet, F. 2008. Distribution of specialist and generalist species along spatial gradients of habitat disturbance and fragmentation. - Oikos 117: 507–514.

15. Didan, K., Munoz, A. B., Solano, R., Huete, A., and others 2015. MODIS vegetation index user’s guide (MOD13 series). - Univ. Ariz. Veg. Index Phenol. Lab 35: 2–33.

16. Dvořáková, L., Kuczyński, L., Rivas-Salvador, J. and Reif, J. 2022. Habitat Characteristics Supporting Bird Species Richness in Mid-Field Woodlots. - Front. Environ. Sci. in press.

17. Dvořáková, D., Šipoš, J. and Suchomel, J. 2023. Impact of agricultural landscape structure on the patterns of bird species diversity at a regional scale. - Avian Res. 14: 100147.

18. Edo, M., Entling, M. H. and Rösch, V. 2023. Agroforestry supports high bird diversity in European farmland. - Agron. Sustain. Dev. 44: 1.

19. Erős, T. and Schmera, D. 2010. Spatio-temporal scaling of biodiversity and the species–time relationship in a stream fish assemblage. - Freshw. Biol. 55: 2391–2400.

20. European Union’s Copernicus Land Monitoring Service information 1990. CORINE Land Cover 1990.

21. European Union’s Copernicus Land Monitoring Service information 2000. CORINE Land Cover 2000.

22. European Union’s Copernicus Land Monitoring Service information 2006. CORINE Land Cover 2006.

23. European Union’s Copernicus Land Monitoring Service information 2012. CORINE Land Cover 2012.

24. European Union’s Copernicus Land Monitoring Service information 2018. CORINE Land Cover 2018.

25. Field, R., Hawkins, B. A., Cornell, H. V., Currie, D. J., Diniz–Filho, J. A. F., Guégan, J., Kaufman, D. M., Kerr, J. T., Mittelbach, G. G., Oberdorff, T., O’Brien, E. M. and Turner, J. R. G. 2009. Spatial species–richness gradients across scales: a meta-analysis. - J. Biogeogr. 36: 132–147.

26. Gábor, L., Jetz, W., Zarzo-Arias, A., Winner, K., Yanco, S., Pinkert, S., Marsh, C. J., Rogan, M. S., Mäkinen, J., Rocchini, D., Barták, V., Malavasi, M., Balej, P. and Moudrý, V. 2023. Species distribution models affected by positional uncertainty in species occurrences can still be ecologically interpretable. - Ecography 2023: e06358.

27. Gooriah, L. D. and Chase, J. M. 2020. Sampling effects drive the species–area relationship in lake zooplankton. - Oikos 129: 124–132.

28. Gorelick, N., Hancher, M., Dixon, M., Ilyushchenko, S., Thau, D. and Moore, R. 2017. Google Earth Engine: Planetary-scale geospatial analysis for everyone. - Remote Sens. Environ. 202: 18–27.

29. Gotelli, N. J. and Colwell, R. K. 2001. Quantifying biodiversity: procedures and pitfalls in the measurement and comparison of species richness. - Ecol. Lett. 4: 379–391.

30. Greenwell, B. M. and Boehmke, B. C. 2020. Variable Importance Plots—An Introduction to the vip Package. - R J. 12: 343–366.

31. Guerrero-Ramírez, N. R., Craven, D., Reich, P. B., Ewel, J. J., Isbell, F., Koricheva, J., Parrotta, J. A., Auge, H., Erickson, H. E., Forrester, D. I., Hector, A., Joshi, J., Montagnini, F., Palmborg, C., Piotto, D., Potvin, C., Roscher, C., van Ruijven, J., Tilman, D., Wilsey, B. and Eisenhauer, N. 2017. Diversity-dependent temporal divergence of ecosystem functioning in experimental ecosystems. - Nat. Ecol. Evol. 1: 1639–1642.

32. Harrison, T., Gibbs, J. and Winfree, R. 2019. Anthropogenic landscapes support fewer rare bee species. - Landsc. Ecol. 34: 967–978.

33. Huntley, B., Collingham, Y. C., Green, R. E., Hilton, G. M., Rahbek, C. and Willis, S. G. 2006. Potential impacts of climatic change upon geographical distributions of birds. - Ibis 148: 8–28.

34. Karger, D. N., Conrad, O., Böhner, J., Kawohl, T., Kreft, H., Soria-Auza, R. W., Zimmermann, N. E., Linder, H. P. and Kessler, M. 2021. Climatologies at high resolution for the earth’s land surface areas.

35. Keil, P. and Chase, J. M. 2019. Global patterns and drivers of tree diversity integrated across a continuum of spatial grains. - Nat. Ecol. Evol. 3: 390–399.

36. Keil, P. and Chase, J. 2022. Interpolation of temporal biodiversity change, loss, and gain across scales: a machine learning approach. in press.

37. Kuhn, M. and Wickham, H. 2020. Tidymodels: a collection of packages for modeling and machine learning using tidyverse principles.

38. Lomolino, M. V., Riddle, B. R. and Whittaker, R. J. 2017. Biogeography. - Oxford University Press.

39. MacArthur, R. H. and Wilson, E. O. 1967. The theory of island biogeography. - Princeton University Press.

40. Mayer, M. 2024. hstats: Interaction Statistics.

41. Miklín, J., Hauck, D., Konvička, O. and Cizek, L. 2017. Veteran trees and saproxylic insects in the floodplains of Lower Morava and Dyje rivers, Czech Republic. - J. Maps 13: 291–299.

42. Moudrý, V. and Šímová, P. 2013. Relative importance of climate, topography, and habitats for breeding wetland birds with different latitudinal distributions in the Czech Republic. - Appl. Geogr. 44: 165–171.

43. Nature Conservation Agency of the Czech Republic 2025. Provision of Data Protected Areas.

44. Prajzlerová, D., Barták, V., Keil, P., Moudrý, V., Zikmundová, M., Balej, P., Leroy, F., Rocchini, D., Perrone, M., Malavasi, M. and Šímová, P. 2024. The relationship between remotely-sensed spectral heterogeneity and bird diversity is modulated by landscape type. - Int. J. Appl. Earth Obs. Geoinformation 128: 103763.

45. Preston, F. W. 1960. Time and Space and the Variation of Species. - Ecology 41: 612–627.

46. Pšeničková, T. and Horák, J. 2022. Influence of forest landscape on birds associated with lowland water bodies. - For. Ecol. Manag. 513: 120199.

47. Pyšek, P., Kučera, T. and Jarošík, V. 2002. Plant species richness of nature reserves: the interplay of area, climate and habitat in a central European landscape. - Glob. Ecol. Biogeogr. 11: 279–289.

48. Rahbek, C. 2005. The role of spatial scale and the perception of large-scale species-richness patterns. - Ecol. Lett. 8: 224–239.

49. Reif, J., Prylová, K., Šizling, A. L., Vermouzek, Z., Šťastný, K. and Bejček, V. 2013. Changes in bird community composition in the Czech Republic from 1982 to 2004: increasing biotic homogenization, impacts of warming climate, but no trend in species richness.- J. Ornithol. 154: 359–370.

50. Ricklefs, R. E. 2004. A comprehensive framework for global patterns in biodiversity. - Ecol.Lett. 7: 1–15.

51. Roberts, D. R., Bahn, V., Ciuti, S., Boyce, M. S., Elith, J., Guillera-Arroita, G., Hauenstein, S., Lahoz-Monfort, J. J., Schröder, B., Thuiller, W., Warton, D. I., Wintle, B. A., Hartig, F. and Dormann, C. F. 2017. Cross-validation strategies for data with temporal, spatial, hierarchical, or phylogenetic structure. - Ecography 40: 913–929.

52. Sandel, B. 2015. Towards a taxonomy of spatial scale-dependence. - Ecography 38: 358–369.

53. Spake, R., Bowler, D. E., Callaghan, C. T., Blowes, S. A., Doncaster, C. P., Antão, L. H., Nakagawa, S., McElreath, R. and Chase, J. M. 2023. Understanding ‘it depends’ in ecology: a guide to hypothesising, visualising and interpreting statistical interactions. - Biol. Rev. 98: 983–1002.

54. Šťastný, K., Bejček, V. and Hudec, K. 1997. Atlas hnízdního rozšíření ptáků v České republice 1985-1989. [Atlas of breeding bird distribution in the Czech Republic 1985-1989]. - H & H.

55. Šťastný, K., Bejček, V. and Hudec, K. 2006. Atlas hnízdního rozšíření ptáků v České republice: 2001-2003. [Atlas of breeding bird distribution in the Czech Republic: 2001-2003]. - Aventinum.

56. Šťastný, K., Bejček, V., Mikuláš, I. and Telenský, T. 2021. Atlas hnízdního rozšíření ptáků v České republice 2014-2017. [Atlas of distributions of breeding birds of the Czech Republic 2014-2017]. - Aventinum.

57. Storch, D. 2016. The theory of the nested species–area relationship: geometric foundations of biodiversity scaling. - J. Veg. Sci. 27: 880–891.

58. Storch, D., Konvicka, M., Benes, J., Martinková, J. and Gaston, K. J. 2003. Distribution patterns in butterflies and birds of the Czech Republic: separating effects of habitat and geographical position. - J. Biogeogr. 30: 1195–1205.

59. Storch, D., Marquet, P. and Brown, J. 2007. Scaling Biodiversity. - Cambridge University Press.

60. Storch, D., Koleček, J., Keil, P., Vermouzek, Z., Voříšek, P. and Reif, J. 2023. Decomposing trends in bird populations: Climate, life histories and habitat affect different aspects of population change. - Divers. Distrib. 29: 572–585.

61. Swenson, N. G., Mi, X., Kress, W. J., Thompson, J., Uriarte, M. and Zimmerman, J. K. 2013. Species-time-area and phylogenetic-time-area relationships in tropical tree communities. - Ecol. Evol. 3: 1173–1183.

62. Tschernosterová, K., Trávníčková, E., Grattarola, F., Rosse, C. and Keil, P. 2023. SPARSE 1.0: a template for databases of species inventories, with an open example of Czech birds. - Biodivers. Data J. 11: e108731.

63. Viana, D. S., Keil, P. and Jeliazkov, A. 2022. Disentangling spatial and environmental effects: Flexible methods for community ecology and macroecology. - Ecosphere 13: e4028.

64. White, E. P. 2004. Two-phase species–time relationships in North American land birds. - Ecol. Lett. 7: 329–336.

65. White, E. P., Adler, P. B., Lauenroth, W. K., Gill, R. A., Greenberg, D., Kaufman, D. M., Rassweiler, A., Rusak, J. A., Smith, M. D., Steinbeck, J. R., Waide, R. B. and Yao, J. 2006. A comparison of the species–time relationship across ecosystems and taxonomic groups. - Oikos 112: 185–195.

66. White, H. J., Montgomery, W. I., Pakeman, R. J. and Lennon, J. J. 2018. Spatiotemporal scaling of plant species richness and functional diversity in a temperate semi-natural grassland. - Ecography 41: 845–856.

67. Wiens, J. A. 2000. Ecological heterogeneity: an ontogeny of concepts and approaches. - Ecol. Consequences Environ. Heterog. 2: 9–31.

68. Williams, C. B. 1943. Area and Number of Species. - Nature 152: 264–267.

69. Wright, M. N. and Ziegler, A. 2017. ranger: A Fast Implementation of Random Forests for High Dimensional Data in C++ and R. - J. Stat. Softw. 77: 1–17.

70. Xie, S., Marzluff, J. M., Su, Y., Wang, Y., Meng, N., Wu, T., Gong, C., Lu, F., Xian, C., Zhang, Y. and Ouyang, Z. 2022. The role of urban waterbodies in maintaining bird species diversity within built area of Beijing. - Sci. Total Environ. 806: 150430.

